# Identifying the therapeutic potential of Niclosamide in overcoming IFN-gamma dependent cancer immune evasion in the Tumor Microenvironment

**DOI:** 10.1101/2025.10.24.684401

**Authors:** Yue Zhang, Sophia Peng, Hamid Akhbariyoon, Suzi Kim, Ellie Lai, En Cai

**Affiliations:** Department of Biological Sciences, Carnegie Mellon University, 5000 Forbes Ave, Pittsburgh, PA 15213; Department of Immunobiology, Yale University, New Haven, CT 06520

**Keywords:** Tumor microenvironment, IFN-γ, PD-L1, STAT1, STAT3, Cancer Stem Cells, Niclosamide, tumor spheroids, Immune Evasion, Cancer Immunotherapy

## Abstract

**Background:** Tumor cells frequently develop immune resistance through interferon-γ (IFN-γ)–induced PD-L1 expression, acquisition of cancer stem cell (CSC)–like features, and adaptation to hypoxia within the tumor microenvironment (TME). Although IFN-γ activates both STAT1 and STAT3, how these pathways interact to regulate immune evasion under hypoxia remains unclear.

**Methods:** Using the MC38 murine colorectal cancer model and T cell–tumor spheroid co-culture assays, we examined how IFN-γ signaling through STAT1 and STAT3 influences PD-L1 expression, CSC plasticity, and cytotoxic T cell function under normoxic and hypoxic conditions. Pharmacologic inhibitors and siRNA knockdown were used to dissect pathway function, and Niclosamide, an FDA-approved anthelmintic, was evaluated as a dual STAT1/STAT3 inhibitor.

**Results:** We found that IFN-γ primarily induced PD-L1 through STAT1 activation, while CSC plasticity was associated with STAT3 signaling. STAT1 and STAT3 displayed reciprocal regulation—blocking one enhanced activation of the other. Niclosamide effectively inhibited phosphorylation of both STAT1 and STAT3, which led to suppressed PD-L1 upregulation and reduced CSC enrichment. In addition, it also partially inhibited hypoxia-induced HIF-1α expression. In co-culture assays, Niclosamide improved T cell infiltration and reduced exhaustion under hypoxic conditions, resulting in improved T cell killing.

**Conclusions:** Our findings identified Niclosamide as a potent dual STAT1/3 inhibitor capable of reversing IFN-γ and hypoxia-driven immune evasion. Repurposing Niclosamide may represent a promising strategy to enhance the efficacy of immune checkpoint blockade in solid tumors.

**key messages:** Interferon-γ (IFN-γ) enhances cytotoxic T cell function but also promotes tumor immune evasion by upregulating PD-L1 and inducing cancer stem cell– like properties. Our study identifies a reciprocal regulatory mechanism between STAT1 and STAT3 in IFN-γ-treated tumor cells that shapes immune evasion outcomes. We demonstrate that Niclosamide, an FDA-approved anthelmintic, acts as a dual STAT1/STAT3 inhibitor, effectively suppressing PD-L1 induction, limiting cancer stemness, and reducing HIF-1α expression under hypoxia. Niclosamide also restores T cell infiltration and decreases exhaustion in a 3D tumor spheroid model. By repurposing Niclosamide, this work provides a feasible approach to enhance the efficacy of immune checkpoint blockade and guide future translational and clinical studies in immunotherapies against solid tumors.

## Background

Over the past decade, cancer immunotherapy, which aims to restore immune function within the tumor microenvironment (TME), has achieved remarkable success in cancer treatment[1,2]. Therapies like immune checkpoint blockade (ICB) have demonstrated durable responses in some patients with advanced malignancies. Nevertheless, only a subset of patients (around 30%) with certain solid tumors benefits from ICBs, while most patients exhibit minimal response to these treatments[3–5]. Understanding the mechanisms of immune evasion and identifying new therapeutic targets remain critical for improving treatment outcomes.

Tumor cells employ multiple strategies to escape immune surveillance. One major mechanism is programmed cell death ligand 1 (PD-L1) upregulation, which inhibits cytotoxic T cell (CTL) function [6,7]. Another mechanism is the development of cancer stem cell (CSC)–like properties, which confer resistance to therapy and drive recurrence[8,9]. In addition, hypoxia is a hallmark of the TME that profoundly reshapes cellular metabolism, promotes PD-L1 expression via hypoxia-inducible factors (HIFs), and suppresses CTL and NK cell functions [10].

During immune responses, T cells release type II interferon (IFN-γ), which is a glycosylated protein that facilitates tumor rejection by modulating systemic anti-tumor immunity[11,12]. However, studies reveal a dichotomous nature of IFN-γ: while it enhances immune functions, it also helps cancer cells evade immune attacks. Specifically, IFN-γ stimulation has been shown to increase cytoplasmic expression of PD-L1[13,14] and elevate cancer cell stemness[15,16]. Both the PD1/PD-L1 axis and CSCs play crucial roles in enabling tumor cells to escape anti-tumor immunity across various cancers[17–19]. IFN-γ activates downstream Janus kinas/signal transducer and activator of transcription (JAK/STAT) signaling, primarily STAT1 and STAT3, which regulate overlapping gene networks involved in immune evasion [20–22]. Therefore, there is an urgent need for therapeutic strategies that inhibit IFN-γ’s effects on cancer immune evasion while preserving its role in facilitating CTL and natural killer (NK) cells in eliminating tumors.

Here, we examined the fine-tuning of the IFN/JAK/STAT pathway under normal and hypoxia condition, and identified the specific STAT pathway that drive PD-L1 upregulation and CSC transformation, respectively. Using a mouse colon cancer model, our study reveals a reciprocal relationship between STAT1 and STAT3 activities in cancer cells (MC38). Specifically, inhibition of STAT1 signaling leads to a significant enhancement in STAT3 activation, and conversely, blocking STAT3 activity results in increased STAT1 signaling. Both transcription factors, STAT1 and STAT3, regulate overlapping sets of target genes, many of which play crucial roles in immune evasion mechanisms. This dynamic interplay suggests that the balance between STAT1 and STAT3 could be a critical factor in shaping the immune microenvironment within tumors, potentially influencing cancer progression and resistance to immune-mediated therapies. Furthermore, in evaluating pharmacological methods to block STAT1/STAT3 activities, we identified an anthelmintic drug, Niclosamide, which inhibits the activity of both STAT1 and STAT3, and partially blocks the upregulation of hypoxia-inducible factor 1α (HIF-1α) protein induced by hypoxia. The co-culture study of primary murine T cells and 3D tumor spheroids model further confirmed that Niclosamide could enhance T cell infiltration and reduce T cell exhaustion under a hypoxic TME. Together, our study reveals Niclosamide as a promising candidate for repurposing in cancer immunotherapy to overcome IFN-γ and hypoxia-induced immune evasion.

## Methods

### Cell Culture and Compounds

MC38 and MC38-OVA cell lines were gifts from Robert Eil lab at OHSU. Both MC38 and MC38-OVA cell line were maintained in DMEM Medium (Gibco, Gaithersburg, MD, USA), supplemented with 10% fetal bovine serum (FBS) and 1% penicillin/streptomycin. Primary OT1 mouse T cells were prepared from lymph nodes and spleens of OT-I TCR transgenic mice (6-10 weeks old), and were maintained in RPMI-1640 Medium (Gibco, Gaithersburg, MD, USA), supplemented with 10% fetal bovine serum (FBS) and 1% penicillin/streptomycin and 50 µ M /J - mercaptoethanol (complete RPMI). Splenocytes were incubated in complete RPMI with 100 ng/mL SIINFEKL peptide for 30 minutes at 37°C and then washed three times. Lymphocytes and splenocytes were then mixed at 1:1 ratio at 2 million cells/mL in complete RPMI and maintained at 37°C 2% CO_2_. Interleukin-2 (HIL2-RO, Roche) were added to the cell culture on day 2 at a final concentration of 10 U/mL. Cells were replenished with fresh media and IL-2 every 2 days and used from day 4 to day 7. Mouse IFN gamma Recombinant Protein were purchased from MilliporeSigma (Sigma-Aldrich, USA). STAT1 inhibitor Fludarabine (F-ara-A, NSC 118218) (MedChemExpress, Cat. No.: HY-B0069), STAT3 inhibitor Stattic (MedChemExpress, Cat. No.: HY-13818), and Niclosamide (BAY2353, MedChemExpress, Cat. No.: HY-B0497) were purchased from MedChemExpress (Monmouth Junction, NJ, USA).

For cell cultured under hypoxia condition, cells were maintained in the Heracell™ VIOS 160i Tri-Gas CO2 Incubator from Thermo Fisher Scientific (Waltham, MA), and nitrogen gas were connected to the incubator to make the oxygen concentration down to 1.5% inside the chamber. For normoxia condition, the oxygen concentration is 20%, as detected by oxygen sensor.

### Mice

C57BL/6-Tg (TcraTcrb)1100Mjb/J (OT1) mice were purchased from Jackson Laboratories (Bar Harbor, ME, USA) and were housed and bred at the Carnegie Mellon University, according to Laboratory Animal Resource Center guidelines. The protocol was approved by the Institutional Animal Care and Use Committee (IACUC) at the Carnegie Mellon University.

### Formation of co-culture tumor spheroids

MC38OVA (or MC38) tumor spheroids were generated by seeding 1×10^4^ cells per well on Costar ultra-low attachment (Corning) round bottom 96 wells plates in the 3D Tumorsphere Medium XF (PromoCell). 7 days later, 1×10^5^ total or CD8 sorted mouse primary T cells were labeled with ViaFluor® 650 SE Cell Proliferation Dye, then added with the spheroids to perform co-culture. Spheroids were gently resuspended and left to sediment to the bottom of the Eppendorf tube. These steps were repeated 2 times with PBS in order to wash the spheroids from the non-infiltrating immune cells. Spheroids were then break down mechanically to obtain a single cell suspension for further analyzed by flow cytometry.

### IFN-γ treatment

5ng/ml IFN-γ treatment for 24 hours were used for most of our experiments, except the experiments with IFN-γ dose and time comparison (Figure 1K, Supplemental Fig.1).

**Figure 1.**
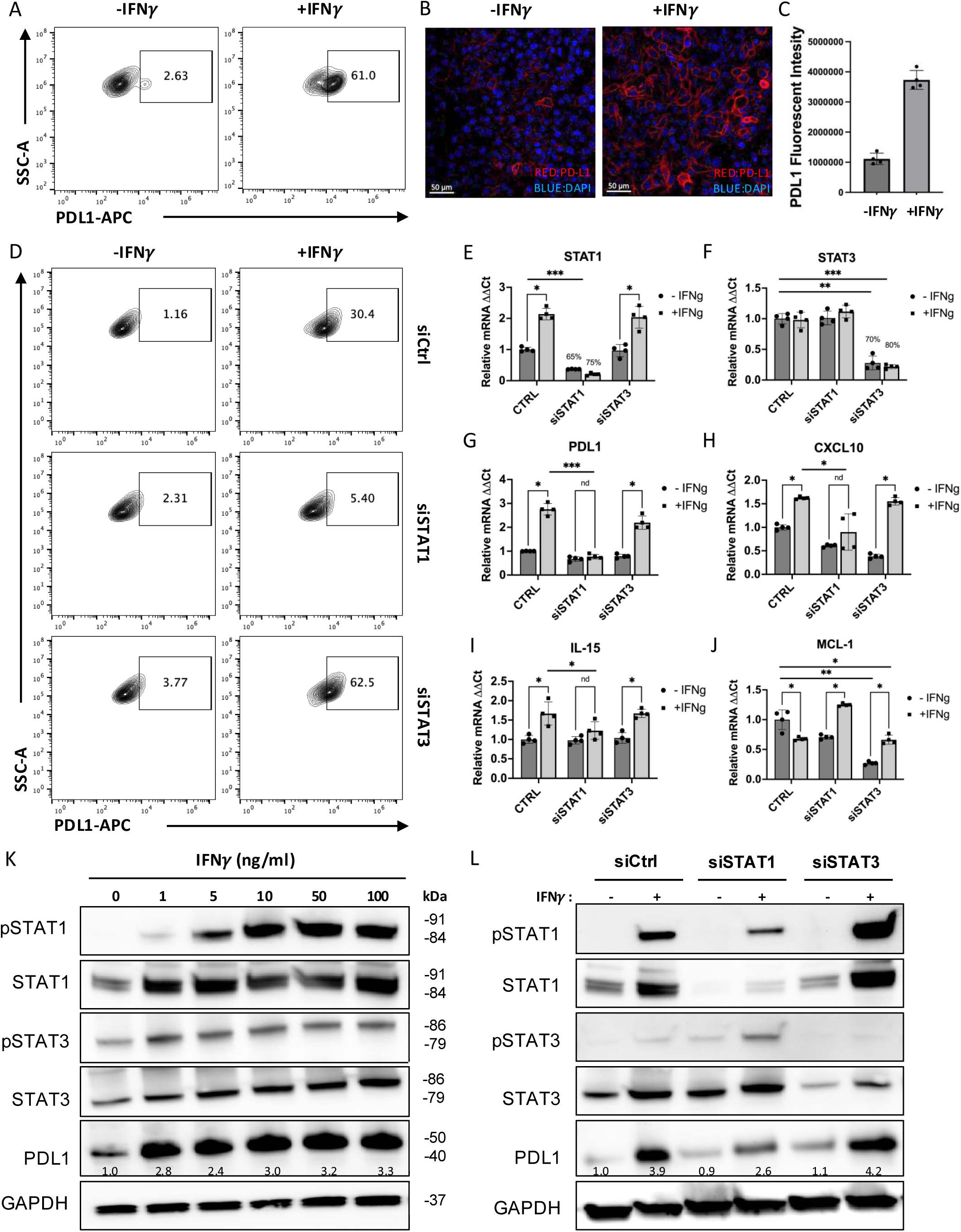
IFNγ up-regulate PD-L1 expression in MC38 cells through IFN-STAT1 signaling. PD-L1 expression on MC38 cell surface was significantly increased in the presence of IFN-γ analyzed by flow cytometry (**A**) and by immune fluorescent staining (**B**-**C**) at 24 hours. (**D**) When treated with siRNA of STAT1, the up-regulation of PD-L1 was significantly reduced, while siRNA of STAT3 has an opposite effect. (**E-F**) The small interfering RNA reaches a sufficient knock down of STAT1 and STAT3, as shown by real-time PCR assay. (**G-I**) The mRNA of STAT1 target genes such as CXCL10 and IL-15 shown similar trend of regulation by IFN-γ as PD-L1. (**J**) The mRNA of STAT3 target gene, MCL-1, shown totally different regulation under normal condition and under treatment of siRNAs. (**K-L)** The expression of STAT1/STAT3 normal and phosphorylated proteins, and PD-L1 proteins in MC38 cell lines were measured by Western blotting with different dose of IFN-γ stimulation **(K**) or after knocking down of STAT1 (siSTAT1) or STAT3 (siSTAT3) **(L**). The number on the image indicates the relative abundance of PD-L1 protein (fold of control). The results are expressed as the mean ± SEM of triplicate measurements in each group. *p<0.05, **p<0.01, ***p<0.001.

### Quantitative real-time PCR

Total RNAs were isolated from cell lysate or mouse tissue with TRIzol reagent (Invitrogen Life Technologies, Carlsbad, CA, USA). The expression levels of multiple mRNAs were detected by quantitative real-time PCR using the High-Capacity cDNA Reverse Transcription Kit (Thermo Fisher Scientific, Waltham, MA, US) and the LightCycler® 480 System (Roche, Indianapolis, IN, USA). Primers used for Quantitative RT-PCR were shown in **Supplemental Table 1**.

### Western Blot

Total proteins were extracted from cultured cells with radio-immunoprecipitation assay (RIPA) buffer (Thermo Fisher Scientific, Waltham, MA, USA) plus fresh protease and phosphatase inhibitors (Roche, Indianapolis, IN, USA). Proteins (10-20μg) were loaded to each lane and separated by sodium dodecyl sulfate polyacrylamide gel electrophoresis.

Proteins were then transferred to a 0.45 µm pore size PDVF membrane (Bio-Rad, Hercules, CA, US), followed by immunodetection of target proteins using specific antibodies with chemiluminescent detection. The developed films were quantified with ImageJ software. The antibodies used in Western blot were listed in **Supplemental Table 2**.

### Flow study

Cell suspensions of MC38 or primary T cells were incubated with the different combinations of antibodies (for MC38: PD-L1, CD133, CD44, B7H4, For T cells: CD45, CD8, PD1, Tim3) on ice for 30 minutes. Then the samples were washed and fixed after cell surface staining according to the manufacturer’s description in fixative buffer(eBioscience). All antibodies were purchased from BD Biosciences and flow data were collected on a BD Accuri C6 Plus Flow Cytometer (BD Biosciences). The antibodies used in flow study were also listed in **Supplemental Table 2**.

The data were analyzed using the FlowJo software (Tree Star Inc.). **Immunofluorescence**: For immunofluorescence, cultured cells or tumor spheres were rinsed with 0.1% TX-100 in PBS (PBST) three times, blocked with 1% goat serum and 2% BSA in PBS at room temperature for 1 h and then incubated with the primary antibody conjugated with fluorescence. Then the slides were washed 3 times with PBST, and mounted with Fluoromount-G™ Mounting Medium (Invitrogen™). Fluorescent images were captured with All-in-One Fluorescence Microscope BZ-X800 (Keyence, Ōsaka, Japan). The primary antibodies used in the immunofluorescence were shown in Supplemental Table 2.

### RNA sequencing (RNA-seq)

RNA-seq of IFN-γ treated MC38 cells was performed by Innomics Inc. (One Broadway, 14th FL, MA) using DNBSEQ Eukaryotic Strand-specific Transcriptome Resequencing. Alignment of raw sequencing reads was performed using Hisat2 software. The DNBSEQ package was used to find differentially expressed genes, and differentially expressed genes with P < 0.05 were used to perform cluster analysis and enrichment analysis with ClusterProfiler package in the R program.

### Statistics

Data were presented as mean ± SEM. Statistical analysis was performed with GraphPad Prism 10.0 software (GraphPad Software, San Diego, CA). Comparison between experimental variables was performed using one-way or two-way ANOVA followed by a Tukey post hoc test. Significance levels were denoted as *p <0.05, **p < 0.01, or ***p <0.001.

## Results

### IFN-γ induced PD-L1 expression through STAT1 pathway

In this study, we used the mouse colon cancer cell line MC38 as a model to study the effects of IFN-γ treatment. Consistent with previous literature reports, we found a significantly increase in PD-L1 expression on MC38 cells following treatment with IFN-γ, as shown by both flow cytometry (Fig.1A) and immunofluorescent staining images (Fig.1B-C). The up-regulation of cell surface PD-L1 demonstrated a dose and time-dependent manner (Supplemental Figure 1). In addition, PD-L1 upregulation was also observed in induced MC38 tumor spheres, which exhibited more CSC features (increased population with CD44+CD133+ expression) compared to adherent MC38 cells (Supplemental Figure 2A-B).

To test how IFN-γ treatment affect tumor cell viability, we used a T cell–tumor cell co-culture system. We found treatment with 5.0 ng/mL IFN-γ enhanced the survival of MC38 cells when co-cultured with primary mouse T cells at a 1:1 ratio. This effect was even more pronounced in tumor spheroids, which exhibited increased resistance to T cell-mediated killing (Supplemental Fig. 2D). Furthermore, MC38 cells pre-treated with 5.0 ng/mL IFN-γ showed markedly increased resistance to T cell cytotoxicity during short-term co-culture (30 min to 4 h) (Supplemental Fig. 3A). Consistently, colon cancer cells exposed to low concentrations of IFN-γ (0.5–5 ng/mL) developed enhanced resistance to T cell killing (at a 1:3 T cell-to-tumor ratio), whereas higher IFN-γ concentrations (10–100 ng/mL) led to decreased viability (Supplemental Fig. 3B). Therefore, we determined to use IFN-γ concentration at 5 ng/mL for our experiments.

Next, we sought to determine the specific STAT signaling pathway that is responsible for IFN-γ induced PD-L1 upregulation. IFN-γ were reported to signal through STAT1 and STAT3 [23,24]. However, the primary signal transducer responsible for IFN-γ dependent PD-L1 upregulation varies between different tumor cell lines[25]. To evaluate the potential roles of STAT1 and STAT3 activation in mediating PD-L1 protein expression in MC38, we used small interfering RNA (siRNA) to knock down STAT1 and STAT3 and examined their distinct contributions to IFN-γ-induced PD-L1 expression. Flow cytometry results showed that IFN-γ-induced PD-L1 expression was diminished when STAT1 was knocked down, while STAT3 knocking-down have an opposite effect (Fig.1D). A similar effect was observed in MC38 cells that were pre-transfected with siRNAs and then induced into tumor spheres (Supplemental Figure 2C). These results suggest that in MC38 cells, IFN-γ upregulate PD-L1 expression through the IFN-STAT1 signaling pathway.

To verify siRNA knock down and assess target gene expression levels, we performed real-time PCR. The results confirmed an efficient knock down of STAT1 (Fig.1E) and STAT3 (Fig.1F) by siRNA. Classical STAT1 transcriptional targets including CXCL10, IL-15 was upregulated in response to IFN-γ treatment, and the up-regulation was diminished with the siRNA of STAT1, but not siSTAT3 (Fig.1H-I). In comparison, myeloid cell leukemia-1 (MCL-1), a transcriptional target of STAT3, was down-regulated by IFN-γ treatment. Knocking down STAT1 increased MCL-1 in the IFN-γ treated sample (Fig. 1J). The mRNA levels of these target genes further confirmed efficient knocking down of STAT1 and STAT3. Notably, when examining PD-L1 levels, we found that IFN-γ-induced PD-L1 upregulation is diminished by knocking down STAT1 but not STAT3. To further assess the total protein expression, we performed western blot, which showed that IFN-γ upregulate PD-L1 in a dose-related manner (Fig. 1K) and that knocking-down of STAT1 with siRNA (siSTAT1) partially blocks the up-regulation of PD-L1 induced by IFN-γ (Fig. 1L). Together, these results confirm that IFN-γ treatment of MC38 cells up-regulate PD-L1 expression through the IFN-STAT1 signaling pathway.

Next, we sought to assess the activation levels of STAT1 and STAT3. We performed western blot assays of the phosphorylated STAT1 (pSTAT1), and phosphorylated STAT3 (pSTAT3) and found that both were enhanced by IFN-γ, and the induced pSTAT1 is much stronger compared to pSTAT3 at higher IFN-γ concentrations. This suggest that IFN-γ activate STAT1 much strong than STAT3. Interestingly, as we compared western blot results of siSTAT1/3 treated samples with scramble siRNA control (siCtrl), we observed that knocking down STAT1 increased pSTAT3 protein level induced by IFN-γ while knocking down STAT3 increased pSTAT1 level induced by IFN-γ (Fig. 1L). This suggests competitive interactions between pSTAT1 and pSTAT3, likely due to the two transcriptional factors compete for similar kinases and promotor sites[25]. This competition explains that knocking-down of STAT3 enhances the increase in PD-L1 expression induced by IFN-γ in western blot (Fig.1L) and flow (Fig.1D). Therefore, IFN-γ-dependent PD-L1 upregulation in MC38 can be suppressed by inhibiting the STAT1 signaling pathway, but this also leads to potential side effects of increased STAT3 activity.

### IFN-**γ** regulate cancer cell stemness indirectly through pSTAT1/pSTAT3 competitive activity

Parallel to our study on IFN-γ-dependent PD-L1 expression in MC38 cells, we seek to investigate the impact of IFN-γ on mitosis and cell cycle. A recent study using mouse models of breast cancer showed that IFN-γ produced by activated T cells can directly convert non-CSCs to CSCs[16]. However, it remains unclear whether this effect is specific to breast cancer models or could be observed across multiple cancer types. Based on the theory that type I interferons (IFNs-I) promote an adaptive yet reversible transcriptional rewiring of cancer cells towards stemness and immune escape[15], we seek to determine how IFN-γ treatment shape the stemness in MC38 cancer cells.

We generated colon cancer spheroids from MC38 cells. We first used ki67 to label the active proliferating cells in mix cell population. To our surprise, we found that IFN-γ treatment reduced the number of ki67 positive cells in a MC38 tumor spheroids model (Fig. 2A-C). To access the stemness of these MC38, we measured the number of sphere-forming unit (SFU) and found it significantly repressed by IFN-γ pre-treatment (Fig. 2D-E), which suggest a reduction in cancer stemness. Furthermore, the reduction of SFU can be neutralized by STAT1 siRNA but not by STAT3 siRNA (Fig. 2F). We next assessed cancer cell stemness with cancer stem cell surface markers, including CD44 and CD133, and observed a decrease of both CD44^hi^CD133^+^ and CD44^hi^CD133^-^ populations in MC38 cells treated with IFN-γ (Fig.2G).

**Figure 2.**
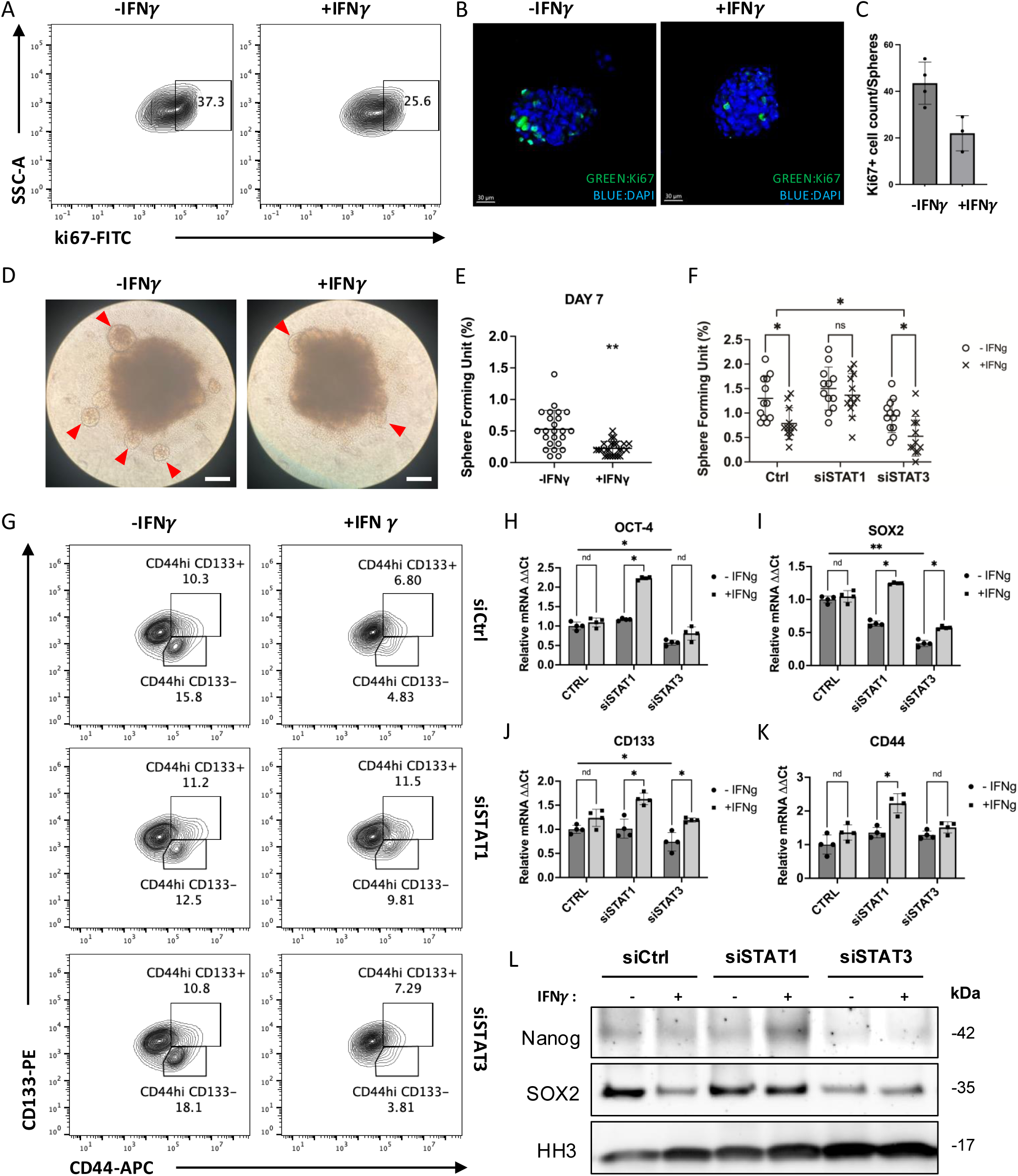
IFNγ modulate cancer cell stemness in MC38 cells indirectly through IFN-STAT3 pathway. (**A-C**) Ki67 expression in the MC38 tumor spheres was significantly reduced in the presence of IFN-γ analyzed by flow cytometry and immune fluorescence. (**D-E**) The tumor sphere forming unit induced in the MC38 cells were also reduced in the presence of IFN-γ (Arrowhead: tumor spheroid. Scale bar: 200μm). **(F)** When MC38 cells were pre-treated with siRNA of STAT1/STAT3, the SFU show different changes. **(G)** When MC38 cells were pre-treated with siRNA of STAT1/STAT3, the CD44+CD133+ population show different trend of regulation by IFN-γ. **(H-K)** The mRNA of cancer stem cell markers was measured by real-time PCR. **(L)** The expression of cancer stem cell marker proteins in cell nucleus was measured by Western blotting. The results are expressed as the mean ± SEM of triplicate measurements in each group. *p<0.05, **p<0.01, ***p<0.001.

Given that these populations were previously reported to represent colorectal cancer stem cells [26–29], our results suggest that CSC stemness decreases upon IFN-γ treatment. Notably, this reduction was abolished only by knocking down STAT1, but not STAT3 (Fig.2G). Accordingly, real-time PCR results demonstrated that IFN-γ can induce CSC markers, including CD44, CD133, OCT4, and SOX2, only when tumor cells were transfected with siSTAT1 (Fig.2H-K). Furthermore, western blot results showed that the protein levels of CSC markers, including SOX2 and Nanog, were reduced upon IFN-γ treatment under normal condition, but remain the same or at higher levels with STAT1 knocking down (Fig.2L). Together, these results showed knocking-down STAT1 resulted in upregulation of CSC-related genes, suggesting that STAT1 playing suppression roles for regulating these CSC-related genes. On the other hand, when MC38 cells were transfected with siSTAT3, both the mRNA and protein of the CSC markers were significantly reduced (except CD44 mRNA, since CD44 is a gene involved in cell–cell interactions, cell adhesion and migration, and may involve other regulation mechanisms)(Fig.2H-L), suggesting that these CSC markers are strongly regulated by STAT3, which is consistent with previously reported [30,31]. The observation that STAT1 knockdown results in upregulation of CSC-related genes, which are known to be strongly regulated by STAT3, provides further evidence supporting the competitive antagonism between pSTAT1 and pSTAT3 as signal transducers and transcriptional factors. Therefore, we conclude that the IFN-γ-induced decrease in CSCs is primarily due to the dominant activation of JAK/STAT1 pathway, which results in increased pSTAT1 that competes with pSTAT3, thereby indirectly suppressing STAT3 signaling. Notably, when STAT1 activity is inhibited, IFN-γ can facilitate the transformation to CSCs through compensatory activation of STAT3 signaling. Therefore, suppressing cancer stemness in the MC38 tumor model requires blocking both STAT1 and STAT3 signaling pathways.

### Niclosamide inhibits both STAT1 and STAT3 activity

We have demonstrated that inhibiting individual STAT1 or STAT3 signaling pathway activated by IFN-γ cannot prevent immune evasion in MC38. To prevent the over-expression of PD-L1 on tumor cells while also suppress the development of CSCs, inhibition of both STAT1 and STAT3 pathways are required.

Next, we sought pharmacological approaches to inhibit IFN-γ induced STAT signaling. We evaluated small molecule STAT1 inhibitor Fludarabine (F-ara-A, NSC 118218) (MedChemExpress, Cat. No.: HY-B0069) and STAT3 inhibitor Niclosamide (BAY2353, MedChemExpress, Cat. No.: HY-B0497). Fludarabine works by inhibiting the cytokine-induced activation of STAT1 and STAT1-dependent gene transcription in normal resting or activated lymphocytes [32,33]. Niclosamide is reported as a STAT3 inhibitor with an IC50 of 0.25 μM in HeLa cells [34]. Both reagents were evaluated in our experiments for blocking STAT1 and STAT3 transcriptional activity respectively.

The western blot result showed that 5μM Fludarabine did not have a sufficient inhibition effect to STAT1 phosphorylation in MC38 cells. In comparison, 0.5μM Niclosamide demonstrated disruption of STAT1 protein expression and phosphorylation while also inhibiting STAT3 protein expression and activity (Fig.3A). In flow-cytometer assays, Niclosamide partially blocked the up-regulation of cell surface PD-L1 induced by IFN-γ (Fig.3B), considering Niclosamide is an STAT3 inhibitor, this effect is opposite compared to STAT3 siRNA (Fig.1D). When low dose of IFN-γ (5ng/ml) were used to increase tumor cell viability in a T cell killing assay, Niclosamide could neutralize this effect while Fludarabine could not (Fig.3C). Meanwhile, Niclosamide did not affect the effectiveness of IFN-γ in reducing SFU (Fig.3D) (Supplemental Figure 4 A-B), nor in reducing the population of CD44^hi^CD133^+^/ CD44^hi^CD133^-^ cell population in MC38 induced tumor spheroids (Fig. 3E).

**Figure 3.**
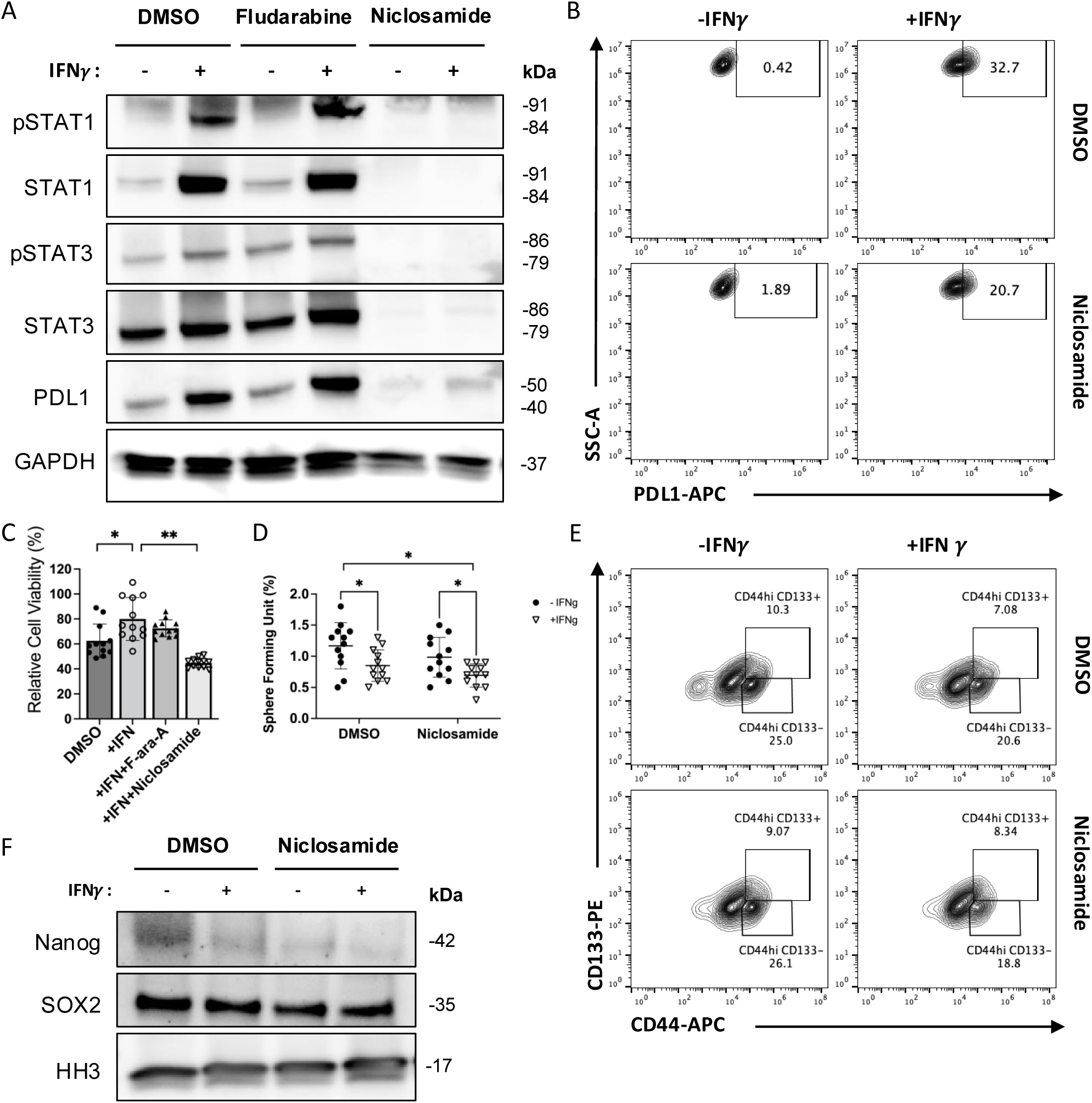
Niclosamide has inhibition effect on both STAT1 and STAT3, which blocks IFNγ-induced PD-L1 up-regulation in MC38 cells while also reduces CSCs formation. **(A)** The expression of STAT1/STAT3 signaling pathway and PD-L1 proteins in MC38 cell lines with Fludarabine and Niclosamide treatment was measured by Western blotting. **(B)** The surface PDL1 expression level was measured by FACS with IFN-γ and Niclosamide. (**C**) The cell viability of MC38 when co-cultured with T cells and pre-treated with Fludarabine and Niclosamide were measured by CCK8 assay. (**D**) The sphere forming units induced in MC38 cells was measured with or without Niclosamide. (**E**) The cell population of CD44+/CD133+ in MC38 treated with Niclosamide was measured by FACS. (**F**) The expression of cancer stem cell marker proteins in cell nucleus was measured by Western blot. The results are expressed as the mean ± SEM of triplicate measurements in each group. *p<0.05, **p<0.01, ***p<0.001.

Additionally, niclosamide reduced PD-L1 expression on the cell surface of these tumor spheroids (Supplemental Figure 4C). In western blot assay, Niclosamide did not affect the downregulation of Nanog, SOX2 proteins by IFN-γ (Fig. 3F). These data suggested Niclosamide could disrupt both STAT1 and STAT3 phosphorylation in IFN-γ signaling, and could become a potential method to prevent cancer cell’s immune evasion in combination use with immune-checkpoint blockers.

### Delineate the signal network in inflammatory TME under hypoxia condition

Hypoxia condition in TME activated a complex signaling network mediated by HIFs, which are closely associated with the JAK/STAT signaling pathways. The hypoxic environment in the solid tumor is well known to mediates resistance to chemotherapy, radiotherapy, and immunotherapy through complex mechanisms[17]. To delineate the changes in multiple signaling pathways during IFN-γ treatment under hypoxia conditions, we sequenced total RNA isolated from the following 4 samples: ***Ctrl_1_1***: MC38 cells under normal condition, without IFN-γ; ***IFN_1_2***: MC38 cells treated with IFN-γ, under normal condition; ***Ctrl_2_1***: MC38 cells under hypoxia condition, without IFN-γ; ***IFN_2_2***: MC38 cells treated with IFN-γ, under hypoxia condition. The total RNA-sequencing analysis provided insights into the transcriptional landscape of cancer cells that has been modulated by the IFN-γ and hypoxia stimulation (Fig. 4A-B). Among all the differentially expressed genes (DEGs), we validate a serious of genes related to cancer progression and immune response with real-time PCR. One of these genes is CD274 (PD-L1), which is significantly up-regulated by IFN-γ alone and further enhanced under hypoxia condition (Fig. 4D). STAT1 and its target genes were all up-regulated by IFN-γ in both conditions (Fig. 4C-F). Notably, the induced CXCL10 (also known as Interferon gamma-induced protein 10, IP-10) level is reduced under hypoxia condition compared to normal condition (Fig. 4E), consistent with literature reports that in certain cancer cells, such as colorectal, breast, ovarian, and lung cancer cells, prolonged hypoxia reduces CXCL10 expression, which is linked to the activity of the HIF-1α protein[35]. According to quantitate PCR results, STAT3 mRNA was also up-regulated by IFN-γ, but its target genes including HIF1α, and some stem cell marker such as SOX2 and OCT-4 mRNA were down-regulated under hypoxia condition (Fig. 4G-N). In general, the RNA-seq results further confirms the role of JAK-STAT signaling in regulating potential genes that mediate cancer immune evasion under hypoxia condition, such as immune-check point blockers (CD274) and chemo attractors (CXCL10).

**Figure 4.**
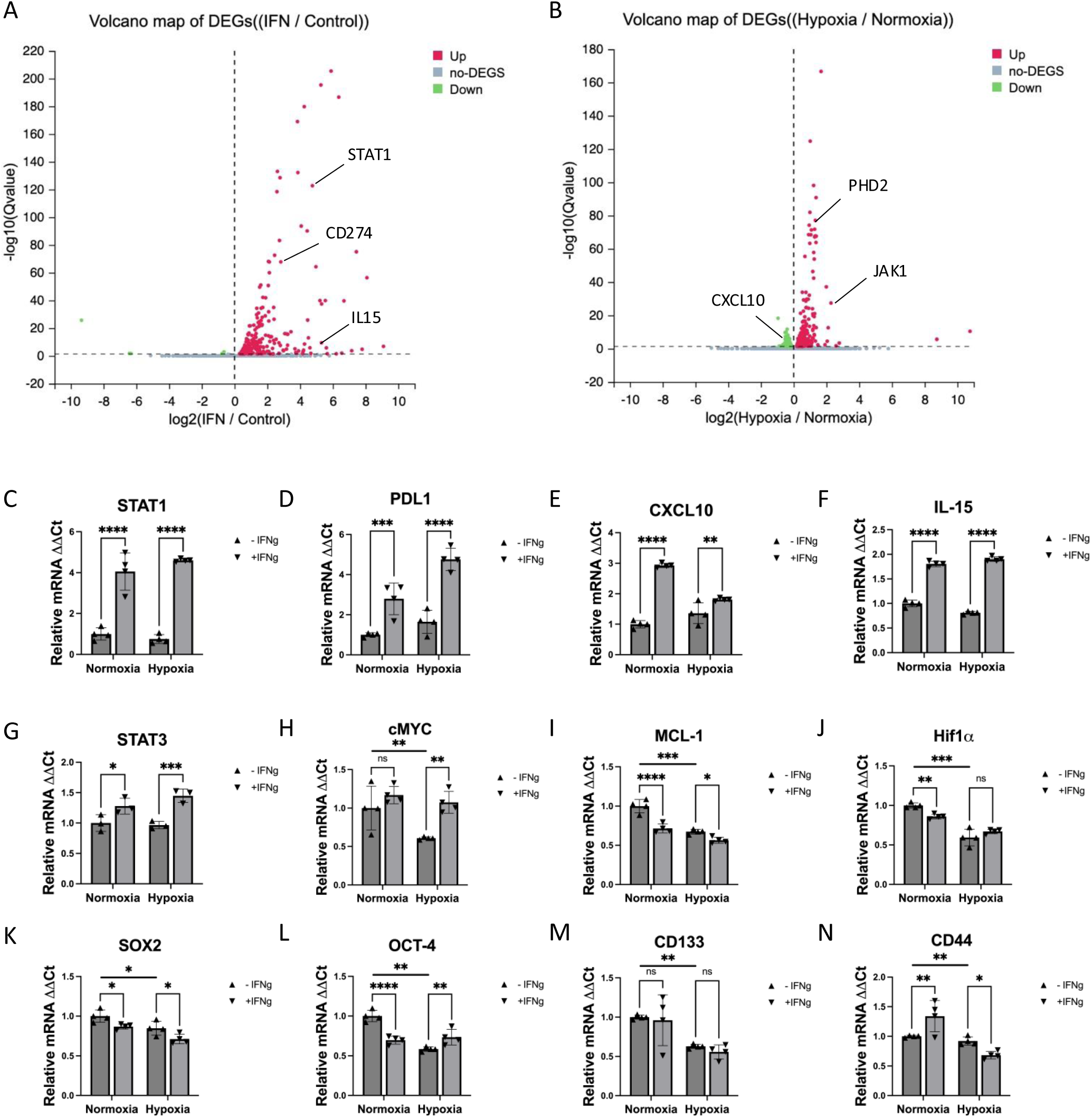
RNA-sequencing analysis from total RNA sample of MC38 cells treated with IFNγ. **(A)** The volcano map of DEGs from RNA-sequencing analysis comparing IFNγ-treated group to control group**. (B)** The volcano map of DEGs from RNA-sequencing analysis comparing hypoxia group to normoxia group. **(C-N)** The real-time PCR validation of varies genes show changes in the RNA-sequencing. The results are expressed as the mean ± SEM of triplicate measurements in each group. *p<0.05, **p<0.01, ***p<0.001.

### Hypoxia enhances IFN-**γ** induced PD-L1 upregulation, which is also blocked by Niclosamide

To further investigate the contribution of hypoxia to immune evasion, we performed western blot analysis of varies protein under both conditions. We found that both PD-L1 and pSTAT1 were up-regulated more significantly in hypoxia condition compared to normal condition (Fig.5A). More importantly, Niclosamide significantly neutralized the induction of HIF1α under hypoxia condition, while preserved its ability to block both STAT1 and STAT3 phosphorylation and protein levels (Fig.5B). The MC38 cells survived better with higher IFN-γ concentration in hypoxia condition compared to normal condition in a T cell-tumor cell co-culture assay (Fig.5C). In a dose-dependent cell viability assay, we found that Niclosamide showed more toxicity to tumor cells under hypoxia condition (Fig.5D). These findings further suggests that Niclosamide could be used as an efficient pharmacological reagent to eliminate IFN-γ-induced immune resistance.

**Figure 5.**
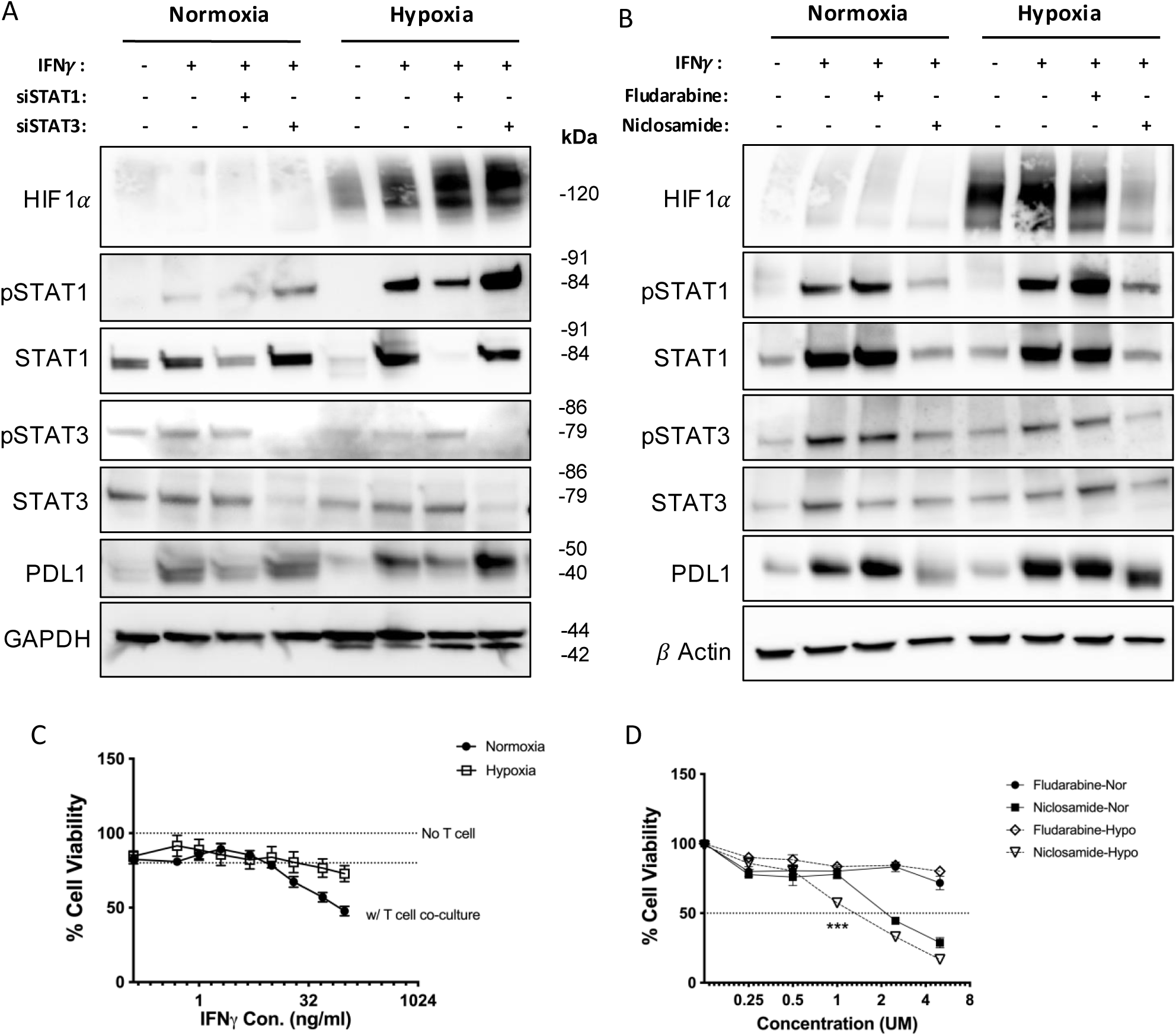
Hypoxia enhances PD-L1 upregulation by IFNγ, while Niclosamide down-regulates Hif1γ under hypoxia condition. **(A-B)** The expression level of STAT1/STAT3 signaling pathway and PD-L1 proteins in MC38 cell lines with siRNA (**A**) or Fludarabine and Niclosamide treatment (**B**) was measured by Western blot. (**C**) Cell viability of MC38 cells pre-treated with different dose of IFN-γ and co-cultured with primary T cells were measured by CCK8 assay, under normoxia or hypoxia condition. (**D**) The Cell viability of MC38 cells treated with different dose of Fludarabine or Niclosamide were measured by CCK8 assay, under normoxia or hypoxia condition. The results are expressed as the mean ± SEM of measurements from 4 ∼ 6 animals per group, and each dot represents a measurement from one animal. *p<0.05, **p<0.01, ***p<0.001.

### IFN-γ induce T cell exhaustion with hypoxia and enhance T cell infiltration in the TME

Given that STATs are also key molecular determinants of T-cell fate and effector function[36], we sought to further study the recruitment and exhaustion progress of T cells in an immunosuppressive tumor microenvironment. We use tumor spheroids induced by MC38-ova cells (transgenic MC38 cell line with stably expressing **Ovalbumin**) and a T cell-tumor cell co-culture system, to mimic a simplified TME[37] (Fig. 6A). Using an established protocol of *in vitro* T cell exhaustion[38], we were able to induce mouse primary CD8+ T cells to an early or late exhausted state (Supplemental Figure 5A). With the exhausted T markers including PD-1, Tim-3, we found that both early state exhaustion (PD-1+/Tim-3-) and late state exhaustion (PD-1+/Tim-3+) T cells were significantly increased when co-cultured with MC38ova cells that were pre-treated with IFN-γ, and under hypoxia condition (Fig. 6B). Meantime, MC38ova cells also showed more induced PD-L1 expression under hypoxia condition (Fig. 6C). We found IFN-γ could promote CD8+ T cells infiltration into the tumor spheres significantly, as shown by CD8+ T cells prelabeled with ViaFluor 650 (VF650) (Fig. 6D-E). We hypothesize this effect is due to CXCL10, a powerful chemokine that acts by binding to its specific receptor, CXCR3, to attract immune cells such as T cells and macrophages to the sites of inflammation. Under hypoxia condition, the T cell infiltration with IFN-γ stimulation is reduced moderately (Fig. 6D-E), which is consistent to our finding that CXCL10 expression is also induced to a lesser degree by IFN-γ under hypoxia condition (Fig. 4E). In addition, ViaFluor® 650 SE dye (**Biotium**, Fremont, CA) is a membrane-permeant compound that is converted to fluorescent dye by intracellular esterases and will covalently reacts with amine groups on intracellular protein at the same time, forming fluorescent conjugates that are retained in the cell. With each cell division, daughter cells inherit roughly half of the fluorescent label, allowing the number of cell divisions that occur after labeling to be detected by the appearance of successively dimmer fluorescent peaks on a flow cytometry histogram. In our experiment, the T cell labeled with viaFluor650 show two peaks in histogram under normal condition, but only one peak under hypoxia condition (Fig. 6F), indicating the T cell proliferation is generally reduced.

**Figure 6.**
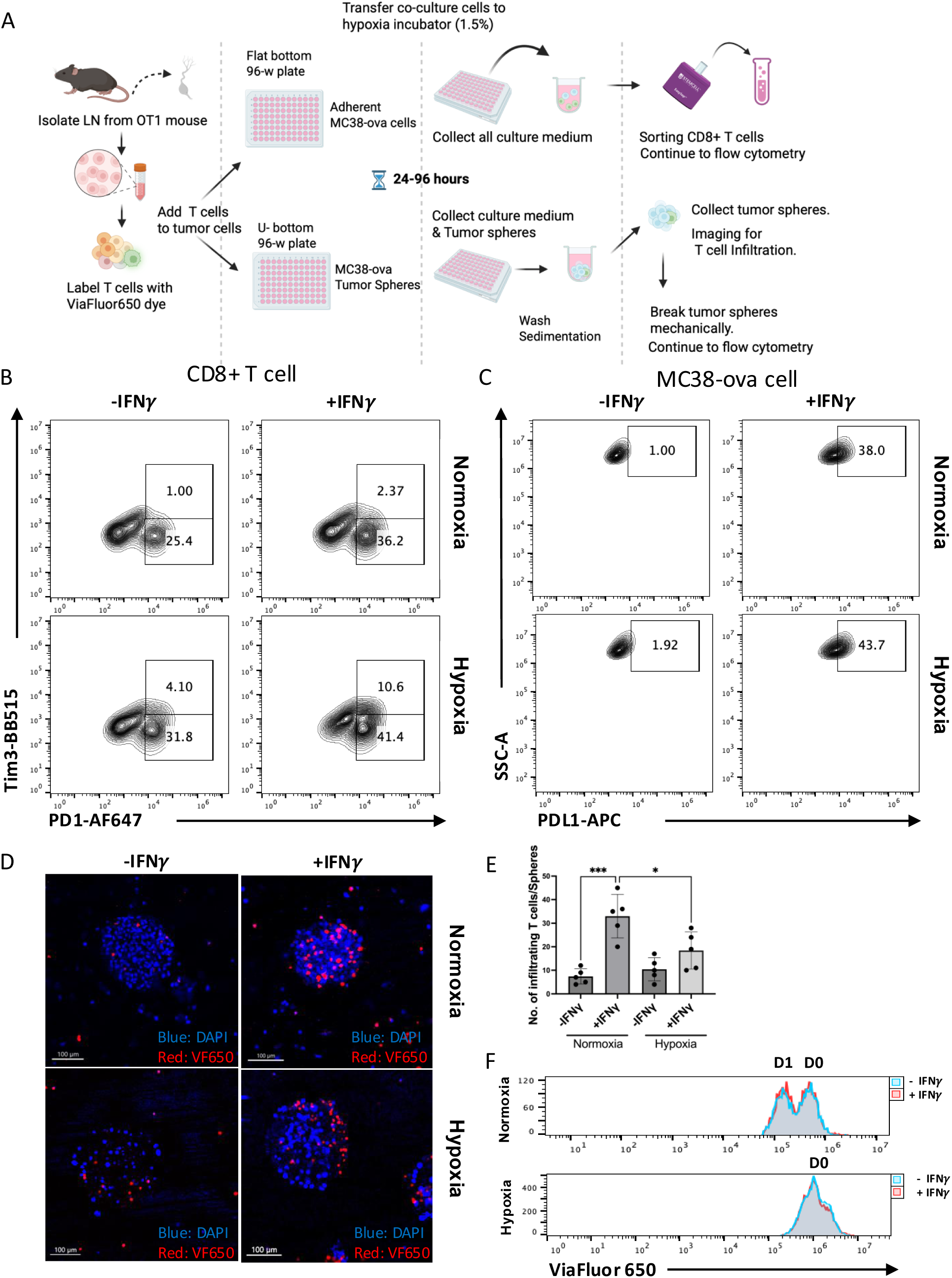
IFNγ and Hypoxia condition has synergistic effect on inducing T cell exhaustion, and IFNγ is essential for T cell infiltration in a tumor-T cells co-culture system. (**A**) Scheme of the protocol used to create co-culture model, using MC38ova adherent cells or tumor-derived spheroids with mouse primary T cells. (**B**) The exhaustion state of CD8+T cells co-cultured with MC38ova cells measured by FACS. (**C**) The expression level of PDL1 on MC38ova cells co-cultured with primary T cells measured by FACS. (**D-E**) The confocal image of tumor-derived spheroids and infiltrating T cells labeled with ViaFluor650 (VF650), with quantification of infiltrating T cell numbers. (**F**) The histogram of ViaFluor650-labeled T cells in the co-culture system measured by flow analysis. The results are expressed as the mean ± SEM of measurements 6 mice per group, and each dot represents a measurement from one animal. *p<0.05, **p<0.01, ***p<0.001.

In another co-culture assay, when the MC38ova cells were pre-treated with both IFN-γ and Niclosamide, the late state exhausted T cell (PD1+Tim3+) population was less compared to IFN-γ treated only condition (Fig.7A). In the T cell-tumor spheres co-culture model, although Niclosamide did not change IFN-γ induced CD8+ T cell infiltration under normal condition (Fig.7B-C), but when IFN-γ’s effect was reduced under hypoxia condition, Niclosamide could partially rescue T cells’ infiltration (Fig.7D). Also, compared to ViaFluor650 labeled CD8+ T cells maintained in media, which divided multiple times in 24hours (D0-D3), the CD8+ T cells get infiltrated into tumor spheroids show only one time division, and IFN-γ increase the subpopulation of divided T cells (D1). Niclosamide did not impair the population of total infiltrated CD8+ T cells (CD8+ Infil), nor the population of proliferated T cells that got infiltrated (CD8+ proli) (Supplemental Figure 5B-C). Together, the inhibition of both STAT1/STAT3 pathway with Niclosamide not only reduces PD-L1 overexpression induced by IFN-γ, but also rescues the hypoxia induced T cell’s exhaustion and less infiltration related to T cell function and metabolism changes. We anticipate that the combination use of IFN-γ plus Niclosamide in colorectal cancer treatment could reach a better therapeutical effect in slowing down the growth of tumors.

**Figure 7.**
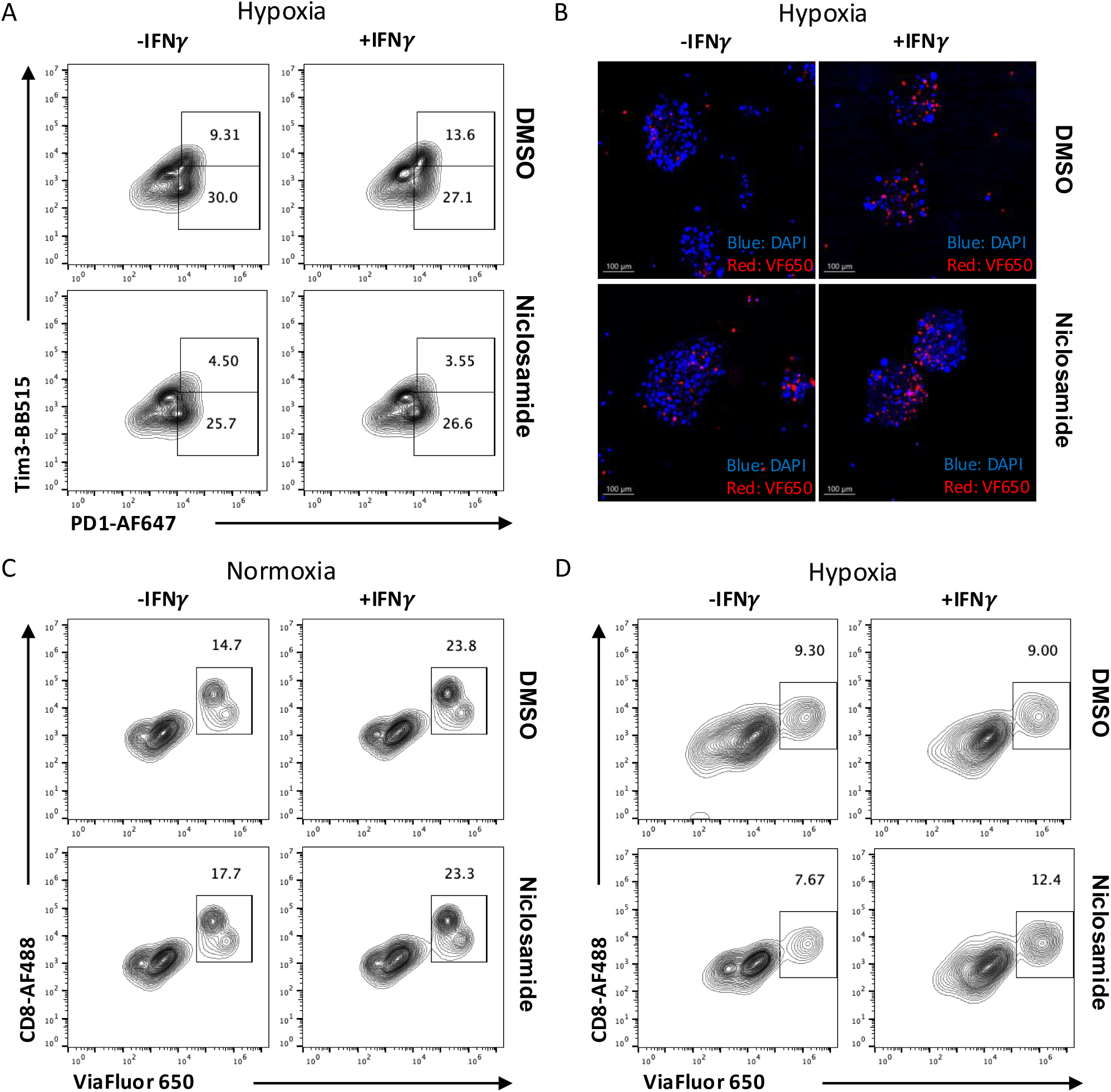
Niclosamide reduces tumor cell immune evasion while preserving IFNγ’s effect in facilitating T cell infiltration in a tumor spheroids-T cells co-culture system under hypoxia condition. **(A)** The exhaustion state of CD8+T cells co-cultured with MC38ova cells measured by FACS, pre-treated with or without Niclosamide. **(B)** The confocal image of tumor-derived spheroids and infiltrating T cells labeled with viaFluor650**. (C-D)** The flow analysis of infiltrating T cells labeled with ViaFluor650 under normal or hypoxia condition. The results are expressed as the mean ± SEM of measurements from 4 animals per group, and each dot represents a measurement from one animal. *p<0.05, **p<0.01, ***p<0.001.

## Discussion

Both hypoxia and consistent inflammation are considered hallmarks of solid tumor development[1,39]. Both conditions drive profound changes in the tumor microenvironment (TME), including metabolic reprogramming, immune suppression, and resistance to therapy [17,18]. Hypoxia impairs immune cell infiltration, promotes regulatory T cells (Tregs) and myeloid-derived suppressor cells (MDSCs), and upregulates PD-L1 expression in tumor and stromal cells [17,18]. These adaptations converge on central signaling hubs such as HIF-1α, STATs, NF-κB, and PI3K-mTOR, which collectively regulate tumor survival and immune evasion [18,40]. Meanwhile, inflammatory cells like tumor-associated macrophages (TAMs) contribute to abnormal, leaky vasculature, exacerbating the hypoxic conditions in tumors[17,41,42]. These findings triggered our great interest since IFN-γ and other cytokines secreted by tumor infiltrated lymphocytes (TILs) also activate the STAT-JAK signaling pathway, and these molecules may have synergistic effect combined with hypoxia condition[21,40].

In this study, we identified a critical interplay between IFN-γ signaling and hypoxia in shaping immune evasion mechanisms. Specifically, we demonstrated that STAT1 mediates IFN-γ-induced PD-L1 expression, while STAT3 promotes cancer stemness, with reciprocal regulation between the two pathways (Fig.1-2). Although IFN-γ could significantly induce PD-L1 up-regulation in the tumor cells, disturbance on the IFN-γ signaling could also result in multiple mechanisms of compensation. Additionally, tumor cells use other strategies to evade immune surveillance and T cells attacking. Importantly, selective inhibition of STAT1 or STAT3 alone enhanced compensatory activation of the other pathway, suggesting that therapeutic strategies targeting only one branch may be insufficient.

We then demonstrated that Niclosamide, an FDA-approved anthelmintic, acts as a dual inhibitor of STAT1 and STAT3 in the MC38 tumor model. Unlike selective inhibitors, Niclosamide simultaneously suppressed PD-L1 upregulation, reduced CSC enrichment, and partially blocked HIF-1α induction under hypoxia (Fig.3, Fig.5). In functional assays, Niclosamide restored T cell cytotoxicity, enhanced infiltration, and reduced exhaustion within a hypoxic TME (Fig.7). Niclosamide is orally bioavailable, inexpensive, and has a well-established safety profile from decades of clinical use against parasitic infections. These properties make it an attractive candidate for rapid translation as an adjuvant to immune checkpoint blockade (ICB). By simultaneously targeting STAT1/3 and hypoxia-driven signaling, Niclosamide may broaden the efficacy of ICB in patient populations that currently exhibit poor response rates. These findings position Niclosamide as a promising repurposed drug candidate for overcoming immune resistance in solid tumors.

Our work also clarifies the paradoxical role of IFN-γ in tumor immunity. While IFN-γ enhances T cell infiltration, it concurrently promotes PD-L1 expression and T cell exhaustion, particularly under hypoxia. This duality highlights the complexity of cytokine signaling in the TME and explains why modulating IFN-γ signaling must be approached with caution. However, our study primarily used *in vitro* and *ex vivo* systems, including MC38 spheroids and T cell co-cultures. While these models capture key features of the TME, *in vivo* validation will be essential to confirm therapeutic efficacy and safety. Moreover, the exact molecular mechanism by which Niclosamide inhibits STAT1 and STAT3 remains to be fully defined and warrants further investigation.

## Conclusion

In general, our study evaluates the effectiveness of targeting STAT1 and STAT3 molecules involved in the IFN-γ pathways to determine which pathway modulator can improve the efficacy of cancer treatments, particularly immunotherapy. Especially, our findings establish Niclosamide as a dual STAT1/3 inhibitor that disrupts IFN-γ-and hypoxia-mediated immune evasion. By restoring T cell function and limiting tumor plasticity, Niclosamide may represent a versatile repurposed agent for combination immunotherapy. This work provides a rationale for preclinical in vivo studies and future clinical trials to evaluate Niclosamide as an adjuvant strategy in refractory solid tumors.

## Acknowledgments

This work was supported by the Shurl and Kay Curci Foundation award to Dr. En Cai. We thank Dr. Eil and Robert Eil lab at School of Medicine, Ohio State University for kindly providing the MC38 and MC38-ova cell lines.

## Author Contributions

Study conception and design: YZ, EC; data collection: YZ, SP, HA, SK, EL; analysis and interpretation of results: YZ, EC; draft manuscript preparation: YZ, EC.

## List of Abbreviations

PD-L1: Programmed Cell Death 1 Ligand 1
CSCs: Cancer Stem Cells
IFN-γ: Interferon-gamma
STAT1: Signal Transducer Activator of Transcription 1
STAT3: Signal Transducer Activator of Transcription 3
JAK: Janus kinase
TME: Tumor Micro-Environment
ICB: Immune Checkpoint Blockade
CTL: Cytotoxic T cells
PD1: Programmed Cell Death 1
HIFs: Hypoxia-inducible Factors
NK: Natural Killer Cells
TAMs: Tumor-associated Macrophages
TILs: Tumor Infiltrated Lymphocytes

