## Supplemental Figures and Tables for "Identifying the therapeutic potential of Niclosamide in overcoming IFN-gamma dependent cancer immune evasion in the Tumor Microenvironment"

**Supplemental Fig. 1 PD-L1 upregulation was observed on the surface of MC38 tumor cells with IFN$\gamma$ treatment of different dosages and time periods.**

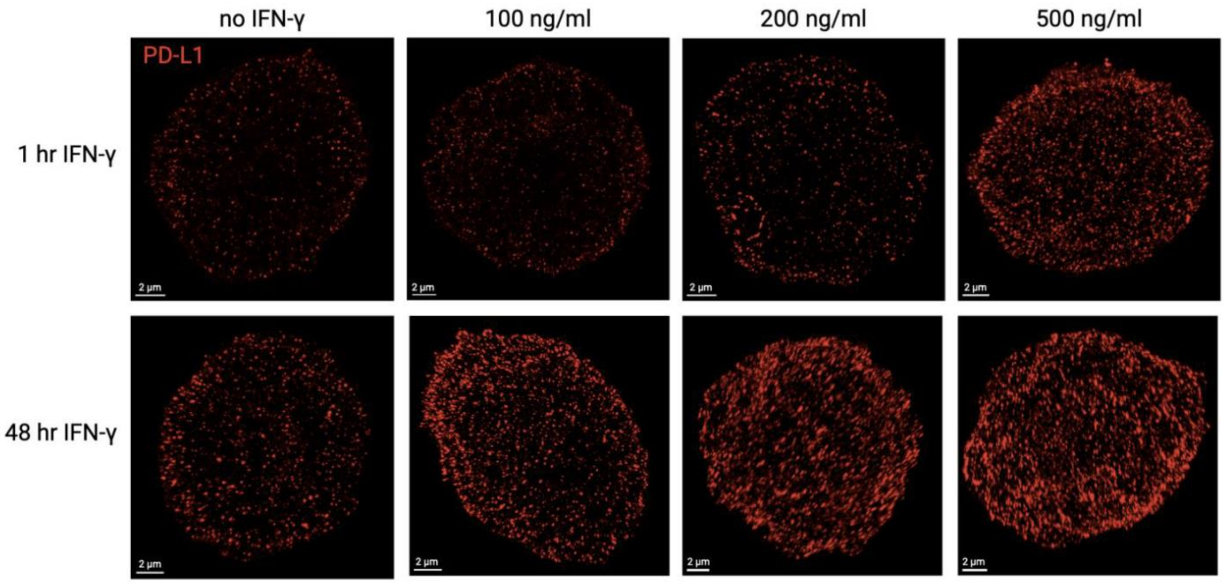


**Supplemental Figure 1. PD-L1 upregulation was observed on the surface of MC38 tumor cells with IFN$\gamma$ treatment of different dosages and time periods.** Confocal image of immune fluorescent staining of surface PDL1 on MC38 cells.

**Supplemental Fig. 2 PD-L1 upregulation by IFN$\gamma$ was observed in induced MC38 tumor spheres, which exhibited more CSC features.**

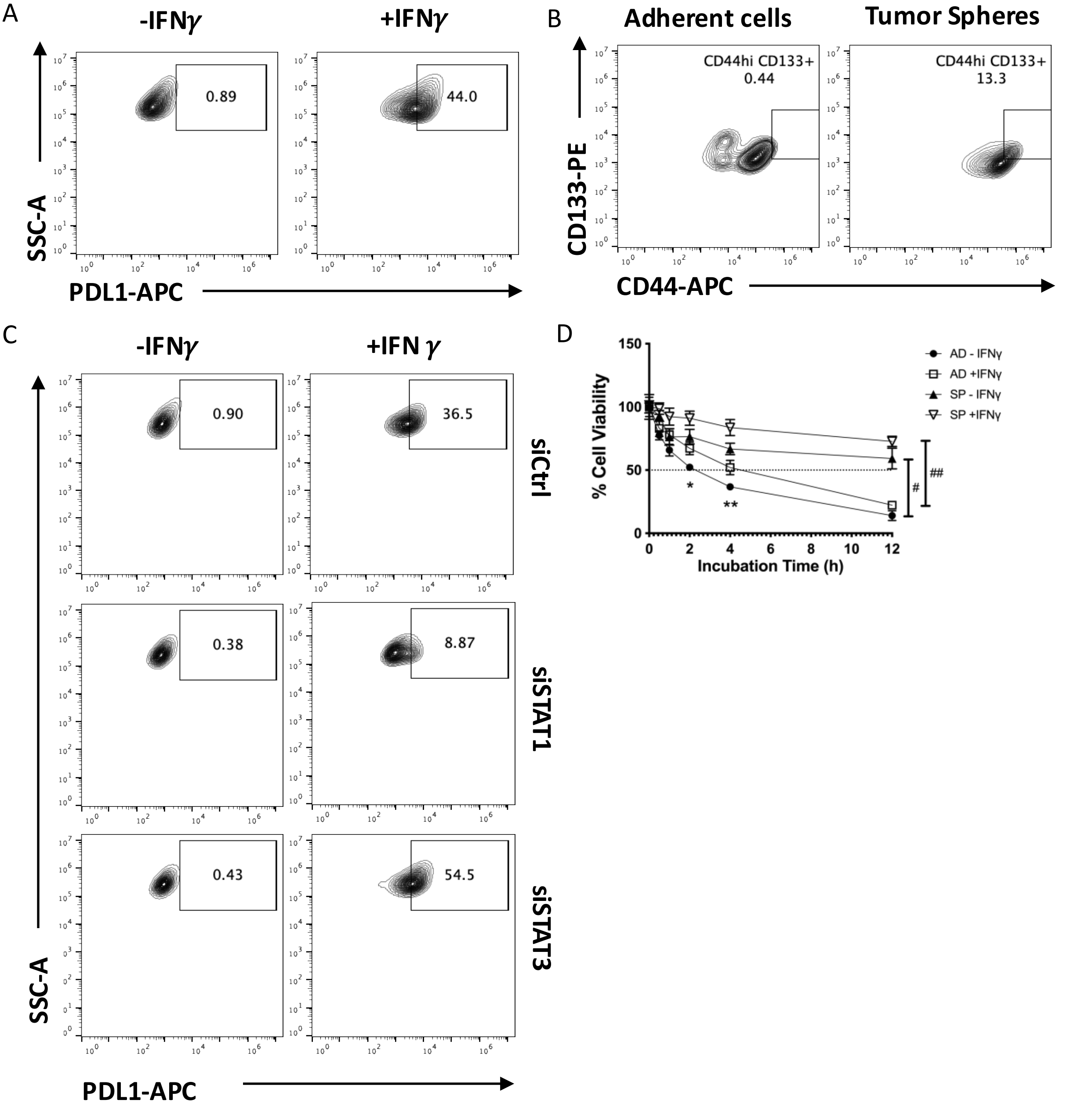


**Supplemental Figure 2. PD-L1 upregulation by IFN$\gamma$ was observed in induced MC38 tumor spheres, which exhibited more CSC features.** (**A**) PD-L1 expression in MC38 tumor spheres level was measured by FACS. (**B**) Flow analysis revealed that the stem cell-like population was much more in tumor spheres compared to adherent cells. (**C**)The PDL1 expression in MC38 cells treated with siRNA of scramble control, STAT1 or STAT3. (**D**) Tumor spheres show more resistant to T cells compared to adherent MC38 cells when co-cultured with T cells. The results are expressed as the mean ± SEM of triplicate measurements in each group. *p<0.05, **p<0.01, ***p<0.001.

**Supplemental Fig. 3 PD-L1 expression induced by IFN$\gamma$ affect tumor cells' viability while co-cultured with primary T cells.**

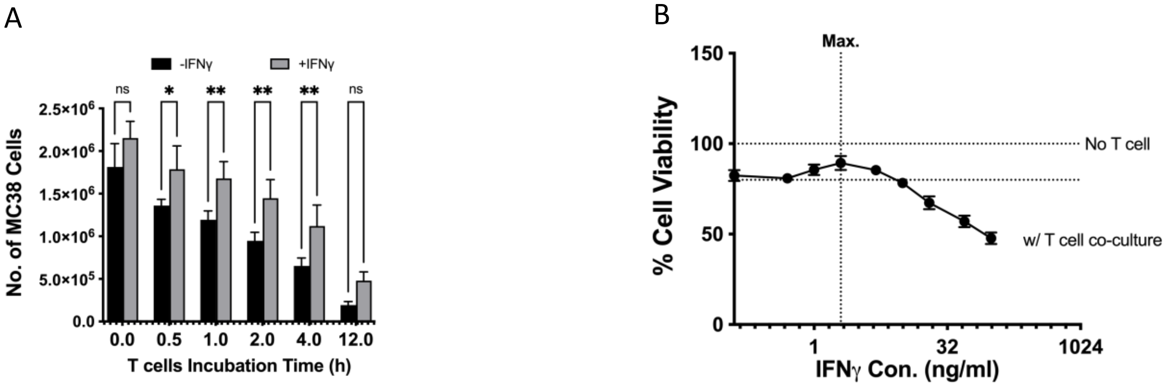


**Supplemental Figure 3. IFN$\gamma$ up-regulate PD-L1 expression affect tumor cells' viability while co-cultured with primary T cells.** (**A**) Live cells number of MC38 cells pre-treated with or without IFNγ and then co-cultured with primary T cells of different time period were measured by Trypan Blue assay. (**B)** Cell viability of MC38 cells pre-treated with different dose of IFNγ and co-cultured with primary T cells were measured by CCK8 assay. *$p<0.05$, **$p<0.01$, ***$p<0.001$.

**Supplemental Fig. 4 Niclosamide reduce tumor spheres formation with or without IFN$\gamma$, while also decrease PDL1 expression in tumor spheres.**

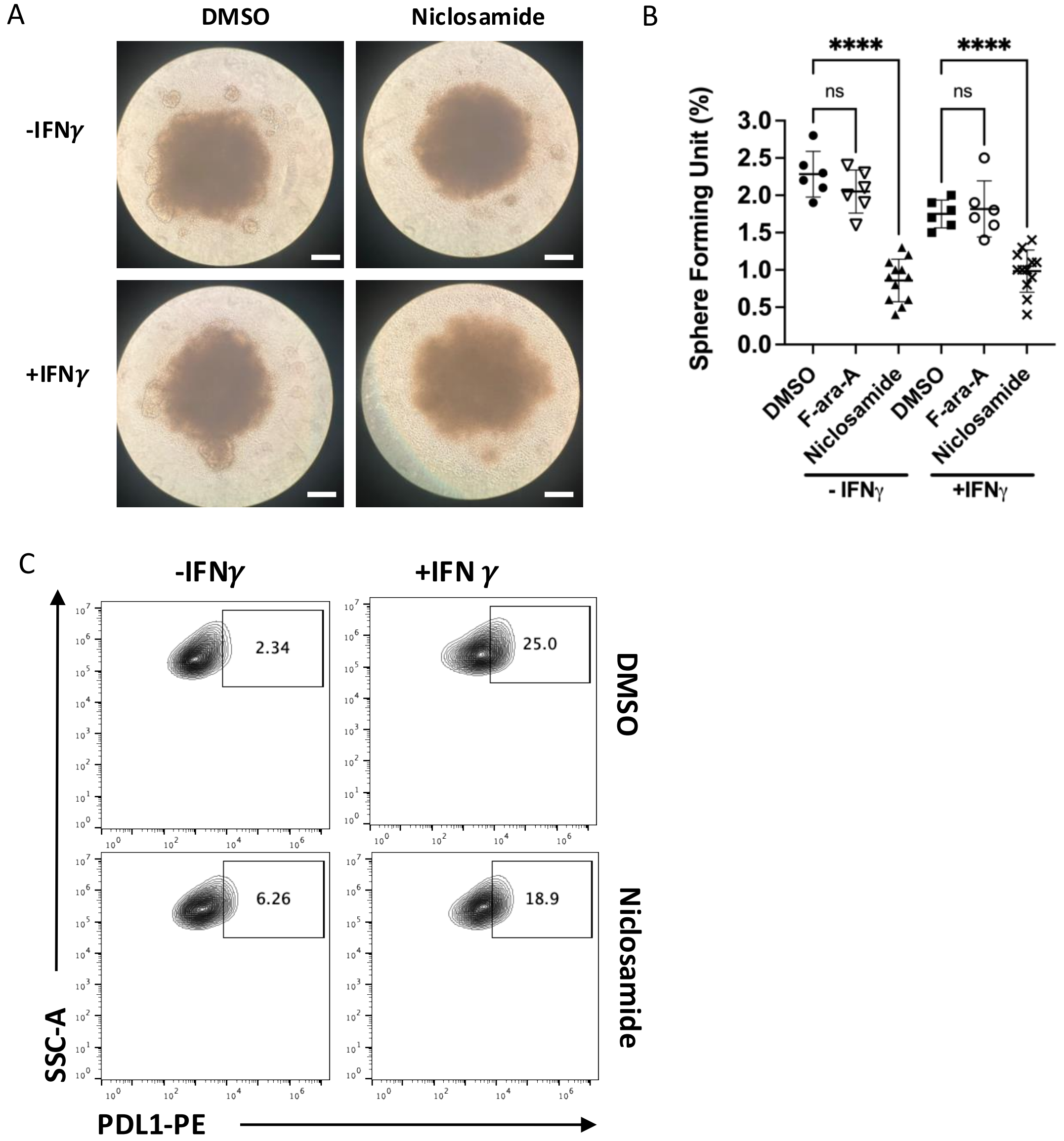


**Supplemental Figure 4. Niclosamide reduce tumor spheres formation with or without IFN$\gamma$.** (**A-B**) The tumor spheres forming unit in the MC38 cells were significantly reduced when treated with Niclosamide, but not with fludarabine (F-ara-A) scale bar: 200μm. (**C**) Niclosamide show partially blocking of the IFN$\gamma$ induced up-regulation of PDL1 in MC38 tumor spheres. The results are expressed as the mean ± SEM of triplicate measurements in each group, ****$p<0.0001$).

**Supplemental Fig. 5 Mouse CD8+ T cells co-cultured with MC38 tumor cells were induced to massive exhaustion under hypoxia condition, and co-cultured with tumor spheroids reduced T cell proliferation.**

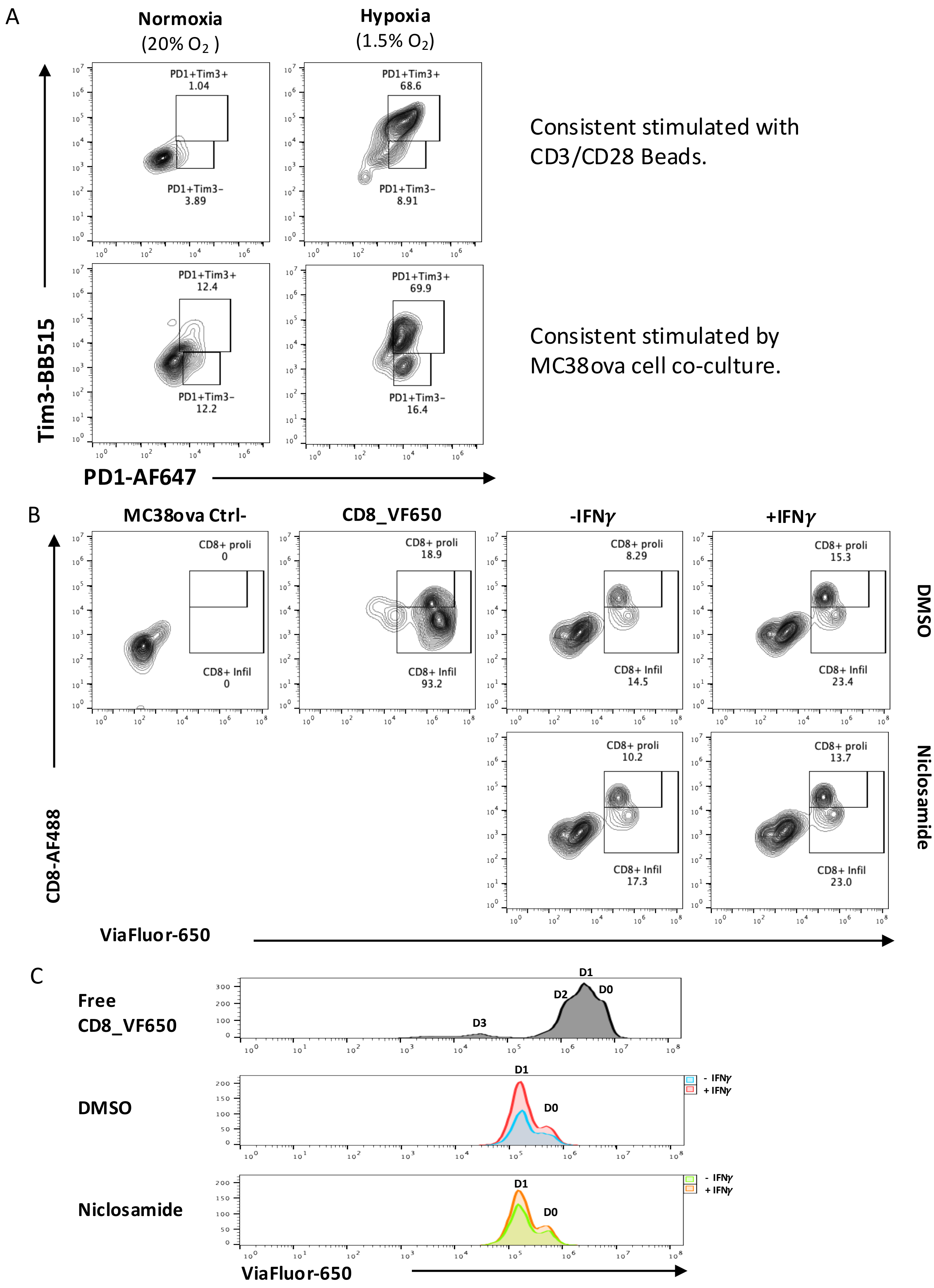

**Supplemental Figure 5. Mouse primary CD8+ T cells co-cultured with MC38 tumor cells were induced to massive exhaustion under hypoxia condition, and co-cultured with tumor spheroids reduced T cell proliferation. (A)** The flow analysis of two different protocol of inducing T cell exhaustion *in vitro*. Upper panel: the primary T cells were co-cultured with CD3/CD28 beads for 7 days. Lower panel: the primary T cells were co-coculture with MC38-OVA cells for 7 days.**(B)** Flow analysis of control MC38ova cells (without staining), CD8+T cells labeled with ViaFluor650, and CD8+ T cells get infiltrated into tumor spheroids, with or without IFN$\gamma$. The gating cell populations are CD8+ T cell get infiltrated into tumor spheres **(CD8+ Infil**), and CD8+ T cells get proliferated after infiltration **(CD8+ proli). (C)** ViaFluor650 dyes are membrane-permeant compounds that are converted to fluorescent dyes by intracellular esterases and will covalently react with amine groups on intracellular proteins at the same time, forming fluorescent conjugates that are retained in the cell. With each cell division, daughter cells inherit roughly half of the fluorescent label, allowing the number of cell divisions that occur after labeling to be detected by the appearance of successively dimmer fluorescent peaks on a flow cytometry histogram. Thus, CD8+ T cell divisions can be tracked after labeling with the ViaFluor650.

### Supplemental Table 1

| Name | Sequence |
|---|---|
| m GAPDH_F | CATCACTGCCACCCAGAAGACTG |
| m GAPDH_R | ATGCCAGTGAGCTTCCCGTTCAG |
| m PDL1_F | TGCGGACTACAAGCGAATCACG |
| m PDL1_R | CTCAGCTTCTGGATAACCCTCG |
| m STAT1_F | GCCTCTCATTGTCACCGAAGAAC |
| m STAT1_R | TGGCTGACGTTGGAGATCACCA |
| m STAT3_F | AGGAGTCTAACAACGGCAGCCT |
| m STAT3_R | GTGGTACACCTCAGTCTCGAAG |
| m SOX2_F | AACGGCAGCTACAGCATGATGC |
| m SOX2_R | CGAGCTGGTCATGGAGTTGTAC |
| m OCT4_F | CAGCAGATCACTCACATCGCCA |
| m OCT4_R | GCCTCATACTCTTCTCGTTGGG |
| m CD44_F | CGGAACCACAGCCTCCTTTCAA |
| m CD44_R | TGCCATCCGTTCTGAAACCACG |
| m CD133_F | CTGCGATAGCATCAGACCAAGC |
| m CD133_R | CTTTTGACGAGGCTCTCCAGATC |
| m Hif1a_R | CCTGCACTGAATCAAGAGGTTGC |
| m Hif1a_F | CCATCAGAAGGACTTGCTGGCT |
| m CXCL10_F | ATCATCCCTGCGAGCCTATCCT |
| m CXCL10_R | GACCTTTTTTGGCTAAACGCTTTC |
| m IL15_F | GTAGGTCTCCCTAAAACAGAGGC |
| m IL15_R | TCCAGGAGAAAGCAGTTCATTGC |
| m cMYC_F | TCGCTGCTGTCCTCCGAGTCC |
| m cMYC_R | GGTTTGCCTCTTCTCCACAGAC |
| m MCL-1_F | AGCTTCATCGAACCATTAGCAGAA |

### Supplemental Table 2

| No | Antibody | Species | Application | Dilution | Manufacturer | Cat. No. |
|---|---|---|---|---|---|---|
| 1 | APC anti-mouse CD274 (B7-H1, PD-L1) Antibody | rat | flow, IF | 1:100 | biolegend | 124312 |
| 2 | FITC anti-mouse/human Ki-67 Antibody | rat | flow, IF | 1:100 | biolegend | 151211 |
| 3 | APC anti-mouse/human CD44 Recombinant Antibody | rat | flow | 1:100 | biolegend | 163603 |
| 4 | PE anti-mouse CD133 Antibody | rat | flow | 1:100 | biolegend | 141203 |
| 5 | BD Horizon™ BB515 Mouse Anti-Mouse CD366 (TIM-3) | rat | flow | 1:100 | BD Biosciences | 567810 |
| 6 | Alexa Fluor® 647 anti-mouse CD279 (PD-1) Antibody | rat | flow | 1:100 | biolegend | 135230 |
| 7 | Alexa Fluor® 488 anti-mouse CD8a Antibody | rat | flow | 1:100 | biolegend | 100723 |
| 8 | APC anti-mouse B7-H4 (B7S1, B7X) Antibody | Armenian Hamster | flow | 1:100 | biolegend | 139407 |
| 9 | PD-L1 (D4H1Z) Rabbit mAb #60475 | Rabbit | WB | 1:1000 | Cell Signaling | 60475T |
| 10 | β-Actin (8H10D10) Mouse mAb #3700 | Mouse | WB | 1:2000 | Cell Signaling | 3700S |
| 11 | GAPDH (D16H11) XP® Rabbit mAb #5174 | Rabbit | WB | 1:2000 | Cell Signaling | 5174S |
| 12 | Phospho-Stat3 (Tyr705) (D3A7) XP® Rabbit mAb #9145 | Rabbit | WB | 1:1000 | Cell Signaling | 9145S |
| 13 | Stat3 (79D7) Rabbit mAb #4904 | Rabbit | WB | 1:1000 | Cell Signaling | 4904S |
| 14 | Phospho-Stat1 (Tyr701) (58D6) Rabbit mAb #9167 | Rabbit | WB | 1:1000 | Cell Signaling | 9167S |
| 15 | Stat1 Antibody #9172 | Rabbit | WB | 1:1000 | Cell Signaling | 9172S |
| 16 | Sox2 (D6D9) XP® Rabbit mAb #3579 | Rabbit | WB | 1:1000 | Cell Signaling | 3579S |
| 17 | Nanog (D2A3) XP® Rabbit mAb #8822 | Rabbit | WB | 1:500 | Cell Signaling | 8822T |
| 18 | Histone H3 (1B1B2) Mouse mAb #14269 | Mouse | WB | 1:500 | Cell Signaling | 14269T |
| 19 | HIF-1α (D1S7W) XP® Rabbit mAb #36169 | Rabbit | WB | 1:500 | Cell Signaling | 36169T |
| 20 | Anti-rabbit IgG, HRP-linked Antibody #7074 | Goat | WB | 1:5000 | Cell Signaling | 7074S |
| 21 | Anti-mouse IgG, HRP-linked Antibody #7076 | Horse | WB | 1:5000 | Cell Signaling | 7076S |
